# Androgen synthesis inhibition but not gonadectomy reduces persistence during strategy set-shifting and reversal learning in male rats

**DOI:** 10.1101/2021.08.11.456005

**Authors:** Ryan J. Tomm, Désirée R. Seib, George V. Kachkovski, Helen R. Schweitzer, Daniel J. Tobiansky, Stan B. Floresco, Kiran K. Soma

**Affiliations:** Department of Psychology and Djavad Mowafaghian Centre for Brain Health, University of British Columbia, Vancouver, BC

**Keywords:** Testosterone, neurosteroids, behavioural flexibility, dopamine, tyrosine hydroxylase

## Abstract

Androgens regulate behavioural flexibility, which is essential to adapt to a changing environment and depends on the medial prefrontal cortex (mPFC). Testosterone (T) administration decreases behavioural flexibility. It is well known that T is produced in the gonads, but T is also produced in the mesocorticolimbic system, which modulates behavioural flexibility. It is unclear how T produced in the brain versus the gonads influences behavioural flexibility. Here, we assess the effects of the androgen synthesis inhibitor abiraterone acetate (ABI) and long-term gonadectomy (GDX) on behavioural flexibility in two paradigms. In Experiment 1, ABI independent of GDX reduced the number of trials to criterion and perseverative errors in a strategy set-shifting task. Similarly, in Experiment 2, ABI but not GDX reduced perseverative errors in a reversal learning task. In subjects from Experiment 1, we also examined tyrosine hydroxylase immunoreactivity (TH-ir), and ABI but not GDX increased TH-ir in the mPFC. Our findings suggest that neurally-produced androgens modulate behavioural flexibility via modification of dopamine signalling in the mesocorticolimbic system. These results suggest novel roles for neurosteroids and possible side effects of ABI treatment for prostate cancer.

## INTRODUCTION

Androgens are well known to regulate reproduction and aggression, but there is increasing evidence that androgens are also critical for more complex behaviours and executive functions, such as behavioural flexibility. In humans, rodents and birds, T treatment reduces behavioural flexibility^1–8^. In these studies, T does not impair learning but rather promotes persistence of a learned strategy and impairs shifting to a novel more advantageous one^9^. The prefrontal cortex (PFC), including the mPFC and orbitofrontal cortex (OFC), is essential for behavioural flexibility^10^. Inactivation of the mPFC and OFC impair set-shifting and reversal learning^11–13^. Importantly, androgen receptors (AR) are expressed in multiple regions of the mesocorticolimbic system, including the mPFC and OFC^14, 15^ as well as the VTA and nucleus accumbens (NAc).

Steroids, including androgens, are locally produced in the brain (neurosteroids) and impact behaviour^16, 17^. Androgens and estrogens are produced in the hippocampus and hypothalamus^18–,22^ and influence learning, reproduction, and aggression^23–26^. The mesocorticolimbic system also produces androgens and estrogens. The mPFC, VTA and NAc express multiple steroidogenic enzymes, including the androgen-synthetic enzyme CYP17A1 and the estrogen-synthetic enzyme CYP19A1 (aromatase)^27, 28^. In these regions, T can act directly via AR or indirectly via aromatization and estrogen receptors (ER). In adult male rats, mesocorticolimbic nodes contain T (measured by mass spectrometry) even after long-term GDX^28^, consistent with local synthesis. Overall, evidence is mounting that the mesocorticolimbic system is androgen-synthetic, but no studies have linked neurally-produced androgens to executive functions such as behavioural flexibility.

Both T and 17-beta-estradiol (E_2_) modulate dopamine transmission, which is essential in regulating behavioural flexibility. These sex steroids can impact dopamine signaling at multiple levels in the mesocorticolimbic system^29^. For example, T affects dopamine production by regulating levels of the dopamine-synthesising enzyme tyrosine hydroxylase (TH) in the mPFC and VTA^30–32^. In addition, T and E_2_ regulate extracellular dopamine levels in the mPFC^33^. Ovarian hormones regulate the number of neurons in the mPFC during puberty^38^. In the NAc, T and E_2_ impact dopamine receptors, transporters, and release^34–37^. Moreover, T affects dendritic spine density in the NAc^39, 40^. Finally, T affects dopamine transporter and dopamine receptor levels in the nigrostriatal pathway^41^. Thus, gonadally- and neurally-produced sex steroids could impact behavioural flexibility via dopamine transmission.

The present study examines the roles of androgens synthesized by the gonads or the brain in regulating behavioural flexibility and dopamine signalling. Adult male rats received long-term GDX and/or androgen synthesis inhibitor ABI^42^. ABI inhibits CYP17A1 activity and crosses the blood-brain barrier^43^. First, to measure behavioural flexibility, we used strategy set-shifting (Experiment 1) and reversal learning (Experiment 2) in separate cohorts^11^. Second, we examined steroid levels in microdissected regions of the mesocorticolimbic system using liquid chromatography tandem mass spectrometry (LC-MS/MS) in subjects from Experiment 1. Third, we measured TH-ir in the mesocorticolimbic system in subjects from Experiment 1. Overall, we hypothesized that GDX would increase behavioural flexibility. We also hypothesized that ABI would increase behavioural flexibility, even in GDX subjects by reducing androgen synthesis in the brain. We also hypothesized that GDX and ABI would affect TH-ir in the mesocorticolimbic system.

## MATERIALS AND METHODS

### Animals

All experiments were authorized by the University of British Columbia Animal Care Committee and performed according to the guidelines of the Canadian Council on Animal Care. Cohorts of 9-week-old adult male Long Evans rats were obtained from Charles River Laboratories (Raleigh, NC/Stone Ridge, NY) weighing 240-300g. We used 2 sets of rats: one set for the strategy set-shifting task (Experiment 1), and a second set for reversal learning (Experiment 2). Animals were group-housed (4 per cage) in clear polycarbonate open top cages (51 cm L x 41 cm W x 21 cm H), and given *ad libitum* access to Rat Diet 5012 (PMI Feeds Inc.) and water for one week upon arrival. Cages contained aspen chip bedding (Nepco), nesting material, and PVC pipes for environmental enrichment. Animals were maintained on a 12 hr light/dark cycle (lights on at 12pm), with the colony room temperature at approximately 21 °C and a relative humidity of 40-50 %.

We designed the experiment as a balanced 2 X 2 factorial design, whereby animals were randomly assigned to 1 of 4 groups. Half of the animals received Sham surgery and the other half received gonadectomy (GDX). Half of each of these 2 groups (Sham and GDX) were then assigned to either Vehicle or ABI treatment producing 4 groups in total: Sham+Vehicle, Sham+ABI, GDX+Vehicle, GDX+ABI. After the surgery and throughout the rest of the experiments, rats were single-housed in clear open-top polycarbonate cages (48 cm L x 27 cm W x 20 cm H), with nesting material and a PVC pipe that were adjacent to one another. Under these conditions, rats can see, smell, and hear conspecifics so they are not isolated per se, and there was no need to control for isolation stress. For 3 weeks, all animals had *ad libitum* access to food and water, after which they were food restricted to 85-90% of their free-feeding weight for the remainder of the experiment (see Figure 1 for experimental timeline), which has often been used to invigorate operant responding.

**Figure 1.**
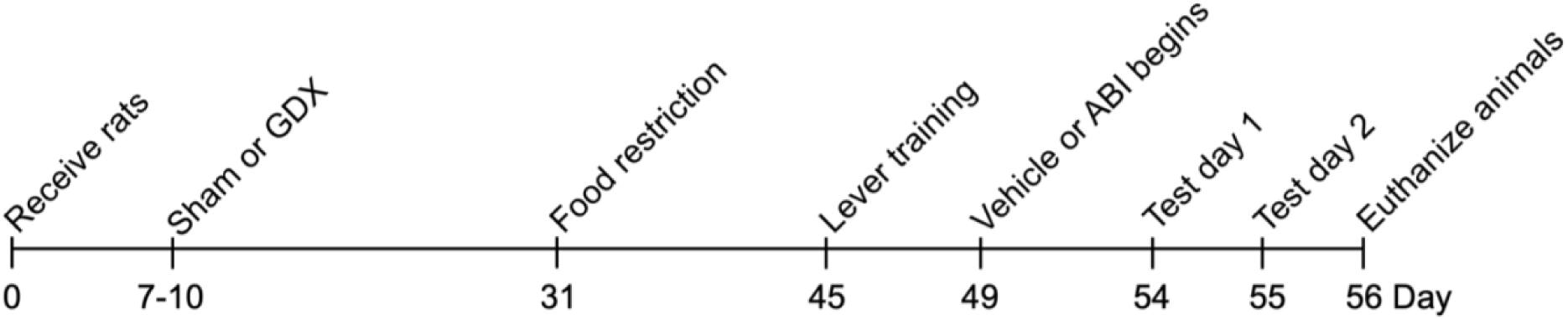
Experimental timeline for Experiment 1 and 2. Male rats were received at 9 weeks of age. 7-10 days after arrival in the facility, rats underwent Sham surgery or GDX. 3 weeks after surgery rats were food restricted and operant training started 2 weeks after food restriction to allow upregulation of local androgen synthesis for a total of 5 weeks. Vehicle and ABI treatment started on day 5 of operant conditioning. Acute tests were performed on day 10 and 11 of operant testing and animals were euthanized and tissue for steroid analysis and immunohistochemistry was collected ∼24h after the last test session. GDX, gonadectomy; ABI, abiraterone acetate.

### Gonadectomy surgery

Surgeries were conducted under aseptic conditions approximately 5 weeks prior to behavioural training as before^28^ to allow for enough time for local T production in the brain to be upregulated^44^ (Figure 1). Briefly, animals were anesthetized with isoflurane and the incision site was numbed with a s.c. bupivacaine injection (2.5mg/ml). For Sham animals, an incision was made to the scrotum and tunic muscle, whilst keeping the vasa deferentia and testes intact. For GDX animals, the procedures were similar, however the vas deferens were bilaterally ligated and the testes removed. Incisions were closed using absorbable sutures, and subcutaneous injections of buprenorphine (0.03 mg/kg) and ketoprofen (5 mg/kg) were administered to alleviate post-surgical discomfort.

### Drug administration

All 4 groups (Sham+Vehicle, Sham+ABI, GDX+Vehicle, GDX+ABI) were habituated for 1 week to 1 g Vehicle, a 50/50 mixture of ground rat chow and peanut butter (Kraft, smooth). The CYP17A1 inhibitor abiraterone acetate (ABI, MedChemExpress, Princeton, NJ; Cat# HY-75054; Batch # 10434) was incorporated in the Vehicle using a food processor for the groups receiving the drug (Sham+ABI, GDX+ABI) so that each 1 g of mixture contained a 40 mg/kg dose of ABI using the average body mass for each group in each cohort. Sham+Vehicle and GDX+Vehicle animals were fed 1 g of Vehicle mixture and Sham+ABI and GDX+ABI animals were fed 1 g of ABI mixture (p.o.) daily starting 5 days prior to behavioural testing and continuing throughout the experiment. To maximize drug effects, Vehicle or ABI was administered 4 hr prior to operant testing when ABI reaches peak levels in tissues after oral administration^43^. A dose of 40mg/kg had been shown to be effective in reducing T levels in serum in rats and mice^45–48^. ABI was given orally since this route is used in humans and has translational value. ABI was given throughout the whole behavioural training and testing phase to allow for genomic effects of T production inhibition.

In a pilot study, male Long Evans rats (∼500-600g) received either Vehicle (n=4) or ABI (n=4, 40mg/kg, p.o.) once per day, for 4 days to test drug efficacy. Animals were habituated for 3 days to vehicle beforehand and euthanized 4 hr after the last ABI administration. The ABI treatment dramatically reduced T levels in serum, mPFC and VTA, as measured by radioimmunoassay^14^ (Drug: F_1,6_ = 9.89, p = 0.02; Tissue: F_1,6_ = 0.82, p = 0.46; Drug x Tissue: F_1,6_ = 0.63, p = 0.55; Table S1). ABI and unconjugated abiraterone were measured by mass spectrometry (details below). We did not detect ABI (conjugated to acetate) but did detect unconjugated abiraterone in serum and brain, indicating robust deconjugation of orally administered ABI. Unconjugated abiraterone was significantly higher in the brain than in the serum (Drug: F_1,6_ = 5.38, p = 0.06; Tissue: F_1,6_ = 6.03, p < 0.05; Drug x Tissue: F_1,6_ = 6.03, p < 0.05; Table S2).

ABI and unconjugated abiraterone were measured using LC-MS/MS. Brain samples (60 - 90 mg) were homogenized in 2 volumes of water and ∼12 ceramic beads using a bead mill (Precellys Tissue Homogenizer; 2 x 20 sec, 6000 rpm). Internal standard (4 µg deuterated T (d3T)) and 10µl 1M HCl were added, and analytes were extracted twice with 1ml of 60/40 hexane/ethyl acetate (hex/EtOAc). Extracts were pooled, dried (CentriVap) and reconstituted in 50 µL methanol. Serum samples (50µl) were similarly processed but without initial homogenization and with 8 µg d3T and reconstituted in 100 µl methanol. Calibration samples were prepared with spiked serum as above to generate a series with 0, 20, 80, 400, 1000 ng/ml ABI/abiraterone. Extracts were analyzed using a Waters Acquity UPLC Separations Module coupled to a Waters Quattro Premier XE Mass Spectrometer. Separations were carried out with a 2.1 x 100 mm BEH 1.7 µM C18 column, mobile phase water (A) and acetonitrile (B) both containing 0.1 % formic acid (gradient: 0.2 min, 40 % B; 2 min, 98 % B; 2-3min, 98 % B; 3.1 min, 40 % B; 5 min run length) at 0.3 ml/min, 35 °C. The MS was set near unit resolution, capillary was 3 kV, source and desolvation temperatures were 120 °C and 300 °C respectively, desolvation and cone gas flows were 1000 L/h and 50 L/h, and the collision cell pressure was held at 4.6 x 10^−3^ mbar. All data were collected in ES+ by multiple reaction monitoring (MRM) with instrument parameters optimized for parent and fragment m/z’s: abiraterone m/z 350.3 > 156.2, m/z 350.3 > 334.3, cone/collision voltages 90/52 and 90/37 respectively; ABI m/z 392.3 > 332.3, cone/collision 50/35; d3T m/z 292 > 97, cone/collision 32/21. Retention times were: abiraterone 1.45 min, ABI 2.4 min, d3T 1.65min. Data processing was done with Quanlynx (Waters) using analyte/IS AUC ratios for quantification and Excel.

### Apparatus

All behavioural testing was done during the light cycle using sound attenuated operant chambers connected to MedPC computer software (30.5 × 24 × 21 cm; Med Associates, St Albans, VT, USA)^11^. Chambers contained a 100mA house light and two retractable levers with a light above each lever. Food reinforcement (1 sugar pellet; 45 mg; Bioserv, Frenchtown, NJ, United States of America) was delivered into a food receptacle located between the 2 levers and infrared photobeams detected locomotor activity.

### Lever press training

Behavioural procedures were adapted from Floresco et al.^11, 49^ for set-shifting and Butts et al.^50^ for reversal learning and performed as described previously^51^. Briefly, the day prior to behavioural training, animals were familiarized with the food reinforcement (∼ 20 sugar pellets) in their home cage. The next day, animals were habituated to the chambers for 30 min, during which food rewards were dispensed intermittently.

Following habituation, animals were trained on a fixed-ratio 1 schedule for each lever (one lever at time; counterbalanced) to a criterion of 60 lever presses. Animals reaching criterion were switched to the opposite lever the next training day. Animals were re-trained on the same lever if criterion was not met. After reaching criteria on both levers, animals were moved to retractable lever press training (90 trials/day) and drug administration began. Here, animals were trained to press an extended lever for food reward. Every 20 sec a trial began with the illumination of the house light and one of the two levers extending. Animals had 10 sec to press the lever for food reward or the trial was counted as an omission and the house light was terminated. After 5 days of retractable lever-press training, animals with fewer than 10 omissions in 90 trials were tested for lever side preference.

### Side preference test

The side preference test was the first time that both levers were presented simultaneously and animals could select between them. The first selection in each trial was tracked over a total of 7 trials and the majority selection was counted as the animals’ side preference. The next day animals began testing on either the set-shifting or reversal learning experiment. For detailed information of the operant training procedures, see Tomm et al.^51^.

### Experiment 1: Strategy set-shifting

#### Visual-cue discrimination

In this experiment 80 male rats were tested (n=19-20 per group). One Sham+Vehicle rat was euthanized prior to behavioural testing and its data were excluded from the analysis. The first test day consisted of 30-150 trials with an inter-trial interval of 20 sec. Every trial began with one of the two stimulus lights (pseudo-randomly, starting either left or right) turning on. After 3 sec, both levers extended and the house light was turned on. To receive a reward, animals had to select the lever with the light above it (i.e., Visual-Cue Discrimination; Figure 2A). If animals did not execute a response within 10 sec, levers retracted, lights extinguished and the trial was counted as an omission. The program was terminated if animals completed 10 consecutive correct choices (after a minimum of 30 trials), or after a maximum of 150 trials.

**Figure 2.**
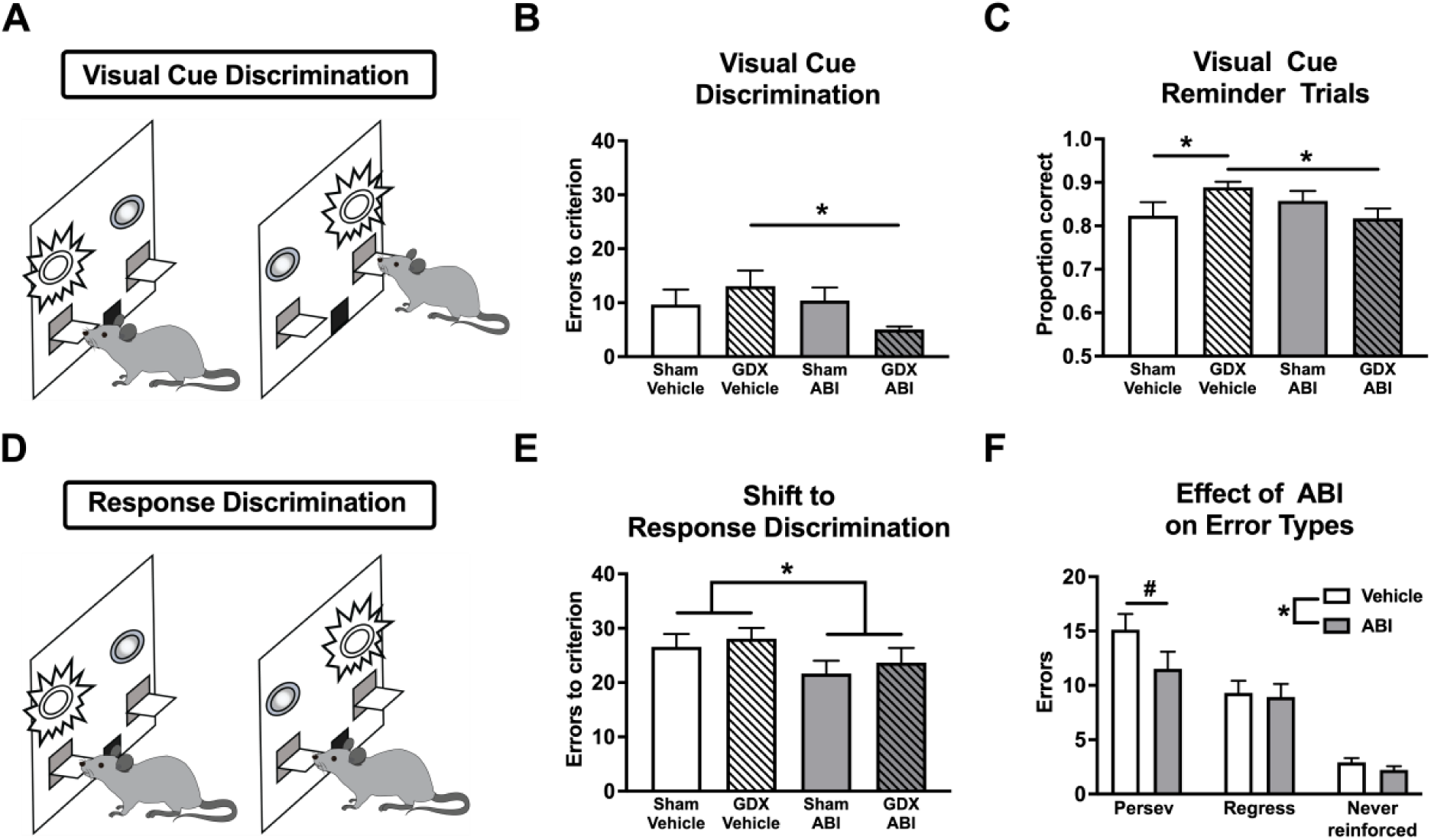
Strategy set-shifting task: Experiment 1. **A** Illustration of the visual cue discrimination. Initially, rats are trained to select the lever with the light above it to receive a reward. **B** Neither GDX alone nor ABI treatment alone affected the errors to criterion during the initial visual cue learning. GDX+ABI resulted in significantly less errors and thus faster learning in this group compared to GDX+Vehicle. **C** GDX+Vehicle animals showed stronger recall of the visual cue discrimination compared to the Sham+Vehicle group as well as the GDX+ABI group. **D** Illustration of the shift-to-response discrimination. Here, rats are required to select one of the two levers (opposite of side bias; e.g. left lever) irrespective of light position to receive a reward. **E** ABI treatment independent of gonadectomy resulted in a reduced number of errors to criterion on the set-shift to the response strategy. **F** Treatment with ABI (Sham+ABI and GDX+ABI pooled) reduced the number of overall errors, which was driven by a reduction in perseverative errors compared to Vehicle treatment (Sham+Vehicle and GDX+Vehicle pooled), indicating a faster shift away from the previously learned strategy or quicker learning of the new strategy. Regressive errors that indicate the ability to maintain the newly learned strategy and never reinforced errors were not significantly affected by ABI treatment. ABI, abiraterone acetate; GDX, gonadectomy; persev, perseverative errors; regress, regressive errors. Data are presented as mean + SEM; n=19-20. *, p ≤ 0.05; #, p = 0.09.

#### Shift-to-response discrimination

The second test day began with 20 trials of the visual-cue discrimination that acted as reminder trials and tested the retention of the initial visual-cue rule. After 20 reminder trials, the rule shifted so animals were now required to always select the lever opposite of the individual animals’ side preference. Thus, animals must disengage from the initial visual-cue rule and shift strategies to respond to an egocentric “response” rule (i.e., always respond to the left lever, irrespective of where the stimulus light is located; Response Discrimination; Figure 2D). The program terminated once the animal completed 10 consecutive correct choices or after 160 trials.

During the set-shift phase, animals could make 3 types of errors: perseverative, regressive, and never-reinforced errors. Perseverative errors were scored when animals responded to the lever with the light illuminated above it on trials that required responding on the opposite lever. Typically, these errors occur in trials immediately following the shift in rules and were scored when 6 or more of this type of error was made in a block of 8 trials. Perseverative errors are indicative of an animals’ inability to disengage from a previously correct but now incorrect strategy^11^. Once animals make 5 or less “perseverative-type” errors in a block of 8 trials, all subsequent errors of this type are scored as regressive errors. Here animals show some understanding of a new strategy, through decreased visual-strategy responding, but still “regress” back to the old visual rule. Regressive errors are indicative of animals’ ability to maintain new strategies. Never-reinforced errors were scored when animals responded to the lever that was neither illuminated with a light, nor the lever opposite to the animals’ side-preference. Here, animals are ignoring both the old visual-strategy and the new response-strategy by making a response that was never reinforced.

### Experiment 2: Spatial reversal learning

#### Response discrimination

For a separate cohort of animals (n=64, n=16 per group), we used the same behavioural training procedures, inter-trial intervals, and omission criterion as the set-shifting task in Experiment 1. The major difference between the reversal learning and the strategy set-shifting task is that, instead of rule shifts between stimulus dimensions (i.e., visual shift-to-response), the rules are reversed within the same stimulus dimension (i.e., response reversal). In this procedure, the visual cue lights still illuminated above one of the levers on each trial, but in this instance, they served as a distractor and did not reliably predict reward. On the first test day, animals had to respond to the lever opposite to their individual side preference, irrespective of light position (randomized in each trial; response discrimination; Figure 5A). Day 1 of the reversal learning task required animals to adopt only an egocentric response-based strategy (opposite to side preference) to reliably receive reward. The program was terminated once animals completed a criterion of 10 consecutive correct choices (after a minimum of 30 trials), or after a maximum of 150 trials. One animal from the GDX+ABI group was excluded from the analysis because of a malfunctioning operant chamber during testing.

#### Response reversal

This test day began with 20 trials of the initial response discrimination that acted as reminder trials and tested the retention of the initial response rule. After 20 trials, the rule was reversed so that animals had to respond to the lever opposite to the initial response rule on Day 1 (response reversal; Figure 5D). During both test days, the lever lights were not predictive of reward. The program terminated if animals completed 10 consecutive correct choices or after 160 trials (after the reversal).

In this experiment, animals could make two types of errors during the response reversal: perseverative and regressive. A perseverative error was scored if animals responded by selecting the lever that was rewarded during the initial response discrimination on Day 1. Perseverative responding was scored when animals made this type of error on 10 or more trials in a block of 16 trials. All other errors were scored as regressive.

### Tissue collection

Only animals from Experiment 1 were used for tissue collection. On the last day of behavioural testing, animals were returned to their home cage and fed. The following day, animals were treated as if they would have another day of testing and received Vehicle or ABI at the usual time of day. However, instead of behavioural testing, animals were euthanized 4 h after drug administration. Animals were transferred one at a time (in their home cage) to a separate room. All animals were rapidly and deeply anesthetized with isoflurane (Pharmaceutical Partners of Canada Inc.) and euthanized either by rapid decapitation (for steroid analysis, n=10 per group) or transcardial perfusion (for immunohistochemistry, n=9-10 per group). Animals were randomly assigned to rapid decapitation or transcardial perfusion conditions. All animals were euthanized in under 3 min (from the start of isoflurane to time of blood collection). This timing is crucial to avoid stress-induced elevation of glucocorticoids^52^.

Animals undergoing rapid decapitation were used to measure steroid levels in whole blood and brain. After rapid decapitation, trunk blood was collected into a microcentrifuge tube. 100 μl of blood were pipetted into another microcentrifuge tube, immediately frozen on dry ice and stored at -80 °C until the time of extraction. Brains were removed, snap frozen in powdered dry ice within 6 min of decapitation and immediately stored at -80 °C.

The other half of animals underwent transcardial perfusion to extract brains for histology. Brains were fixed in 4% PFA for 4h for analysis TH-ir. For detailed experimental procedures of perfusion, see Low et al.^14^.

### Steroid extraction and measurement

Brains from Experiment 1 used for steroid measurement were sectioned coronally in a cryostat (- 12 °C) at 300µm. Sections were then placed on positively charged glass microscope slides for storage (-80 °C) until micro-dissection procedures. Micro-dissection of the brain using the Palkovits punch method^28, 51–53^ allows for collection and measurement of steroids in discrete brain regions^15^. We measured steroid levels in the lateral orbitofrontal cortex (LOFC), the prelimbic region of the mPFC, NAc, dorsomedial striatum (DMS), and the VTA (see Figure S1). We did not measure steroid levels in the medial orbitofrontal cortex (MOFC), since the region was too small to specifically target with our method. Brain regions were located using Paxinos and Watson’s brain atlas^54^, and a Nissl stain was used to verify the locations of punches. One NAc sample in the GDX+ABI group was excluded from our analysis since the brain punches fell outside of their respective coordinates.

Steroid extraction and measurement was adapted from Tobiansky et al.^28^ with minor modifications, and the addition of progesterone to the steroid panel. Briefly, steroids were extracted from brain punches and whole blood (3 µl). Deuterated internal standards T-d5, corticosterone-d8, progesterone-d9, 17β-estradiol-d4, and DHEA-d6 were added. Blanks and standard curves were prepared using 0.2-1,000 pg/tube for T, corticosterone (CORT), progesterone, E_2_, and 4-10,000 pg/tube for dehydroepiandrosterone (DHEA).

Samples were resuspended in 500 µL of HPLC-grade methanol and vortexed for 10 sec. Steroids were extracted as described by Tobiansky et al. and Hamden et al.^28, 55^ via solid phase extraction in C18 columns (Agilent, cat. 12113045). In the last step, samples were resuspended in 50 µL of 50% HPLC-grade methanol in MilliQ water, centrifuged for 2 min at 16,100 g, transferred to glass vial inserts, and steroids were measured with LC-MS/MS as previously described^28^. Steroids were detected with multiple reaction monitoring (MRM), with two MRM transitions for T, CORT, progesterone, E_2_, and DHEA, and one MRM transition for internal standards. Steroid concentrations were acquired on an AB Sciex 6500 Qtrap triple quadrupole tandem mass spectrometer (AB Sciex LLC, Framingham, MA) in a positive electrospray ionization mode for T, CORT, Progesterone, and DHEA, and in a negative electrospray ionization mode for E_2_.

### Tyrosine hydroxylase expression

#### Immunohistochemistry

Immunohistochemistry was performed on animals from Experiment 1 as before^51^. Briefly, after fixation, brains were washed in phosphate buffered saline (PBS) for 1.5 hr (3 x 30 min) before cryo-protection in 30% sucrose for 48 hr at 4 °C. Brains were then removed from solution and stored at -80 °C until sectioning. Brains were sectioned at 40µm coronally in a cryostat at -20 °C. Free-floating sections were placed in cryoprotectant and stored at -20 °C until immunohistochemistry. TH staining procedures were as before using well-plates^8, 14^. Importantly, one subject from each experimental group was stained at the same time in the same plate to facilitate comparisons across groups. Antibodies used were anti-TH (1:2000; clone LNC1, mouse monoclonal; Millipore; Billerica, MA; Cat# MAB318) and secondary goat anti-mouse antibody (1:2000; Biotin-SP-conjugated AffiniPure IgG; Jackson ImmunoResearch Laboratories; West Grove, PA; Cat# 115-065-116). Immunoreactivity was faint in cortical regions, since TH is located mainly in projections and not cell bodies, so another series was used to stain TH in the PFC.

#### Quantification procedures

For the mesolimbic Regions (NAc, DMS, and VTA), we quantified TH-ir using mean greyscale intensity as before^28^. Briefly, experimenters blind to conditions captured photomicrographs using an Olympus CX41RF microscope (10x objective; 2048 x 1532 resolution; Tokyo, Japan) fitted with an Olympus SC30 camera (Tokyo, Japan) and processed images using Fiji ImageJ^57^ (ImageJ 1.34s software; NIH, US). For the NAc, DMS, and VTA, images were taken from 3 sections per region: one rostral, one central, and one caudal image. The regions were located using Paxinos and Watson’s Rat Brain Atlas^54^, and included the NAc shell (NAcS; bregma +2.28 to +0.96), NAc core (NAcC; bregma +2.28 to +0.96), DMS (bregma +1.56 to -0.24) and the parabrachial pigmented nucleus of the VTA (bregma –4.68 to −6.00; Figure S2). The ROIs for each region were formed by cropping 10x images into squares (150 × 150 pixels; 1.32 μm/pixel) and converting the images into 8-bit greyscale format, which assigned each pixel a greyscale value ranging from 1 to 256 (total black to total white). For each image, an average greyscale value was determined, which were compared across groups, as done before^58, 59^.

For the cortical Regions (mPFC and OFC), because of faint staining, we were unable to utilize the mean greyscale method described above. Instead, we used a method in which we compared the amount of staining signal to the background as a percentage of positive pixels. The same procedures for staining, imaging, and software was used as above, and again we used Paxinos and Watson’s Rat Brain Atlas^54^ to locate cortical regions, including the MOFC (anterior-posterior from bregma (AP): +5.60 to 4.20), LOFC (bregma +5.16 to +3.00), and layer 2/3 of the prelimbic mPFC (PrL; bregma +5.16 to +2.76). Again, a total of 3 sections per region were analyzed as above. Images were converted to 8-bit greyscale, inverted, and thresholded using 1.5-fold intensities compared to the background to identify positive staining. The output percentage value representing the amount of stain positive pixels within a selection of the image (600x600 pixels; 1.32 μm/pixel) was compared between groups. Statistical analysis

For strategy set-shifting, the primary dependent variable was the number of errors to criterion during the shift-to-response discrimination. This was analyzed using a 3-way between-/within-subjects ANOVA, with Drug and Surgery as between-subjects factors and Error Type (perseverative, regressive, never-reinforced) as the within-subjects factor. This analysis was followed up with a 1-way ANOVA testing the effects of Drug on each error type. Similarly, for reversal learning, a 3-way between-/within-subjects ANOVA was used for our primary dependent variable looking at the number of errors to criterion during the response reversal. This ANOVA looked at Drug and Surgery as between-subjects factors and Error Type (perseverative, regressive) as the within-subjects factor and was followed up with 1-way ANOVAs looking at the effects of Drug on each error type. All other behavioural analysis, including trials to criterion, omissions, response latencies, and locomotor activity employed 2-way between-subjects ANOVAs with Drug and Surgery as between-subjects factors, and the appropriate analysis of the simple main effects was used when necessary.

For the steroid analysis and histology, only animals from Experiment 1 were analyzed. The primary dependent variable for each study was the concentration of T, CORT and Progesterone in whole blood and the brain. Each steroid was analyzed separately using a 3-way between-/within-subjects ANOVA, with Drug and Surgery as between-subjects factors and Tissue Type (whole blood, LOFC, mPFC, NAc, DMS, VTA) as within-subjects factor. For each steroid, between-/within-subjects ANOVAs, with Drug and Surgery as between-subjects factors, and Tissue Type as within-subjects factors was used, first looking at steroid levels in whole blood and then looking at brain steroid levels (LOFC, mPFC, NAc, DMS, VTA) separately for the within-subjects factor. All statistical analysis was performed using R (Version 4.0.2, “Taking Off Again”)^60^. Data are available upon request from the corresponding author.

## Results

### Experiment 1: Strategy set-shifting

#### Day 1: Visual-cue discrimination

During the visual-cue discrimination learning, there were no main effects of Drug or Surgery on trials to criterion (Drug: F_1,75_ = 1.07; p = 0.30; Surgery: F_1,75_ = 0.44, p = 0.51; Table 1) or errors to criterion (Drug: F_1,75_ = 2.48; p = 0.12; Surgery: F_1,75_ = 0.17, p = 0.68; Figure 2B). There was, however, a significant Drug x Surgery interaction on trials to criterion (F_1,75_ = 5.55, p = 0.02), whereby ABI decreased trials to criterion in GDX rats compared to Sham+ABI rats (F_1,75_ = 4.62, p = 0.03; Table 1). We observed a similar trend on errors to criterion for Drug x Surgery interaction (F_1,75_ = 3.54, p = 0.06). As a result, rats in the GDX+ABI group made the least errors to reach criterion compared to the GDX+Vehicle group (trials: F_1,75_ = 5.83, p = 0.02; Table 1; errors: F_1,75_ = 6.05, p = 0.02; Figure 2B).

**Table 1.**
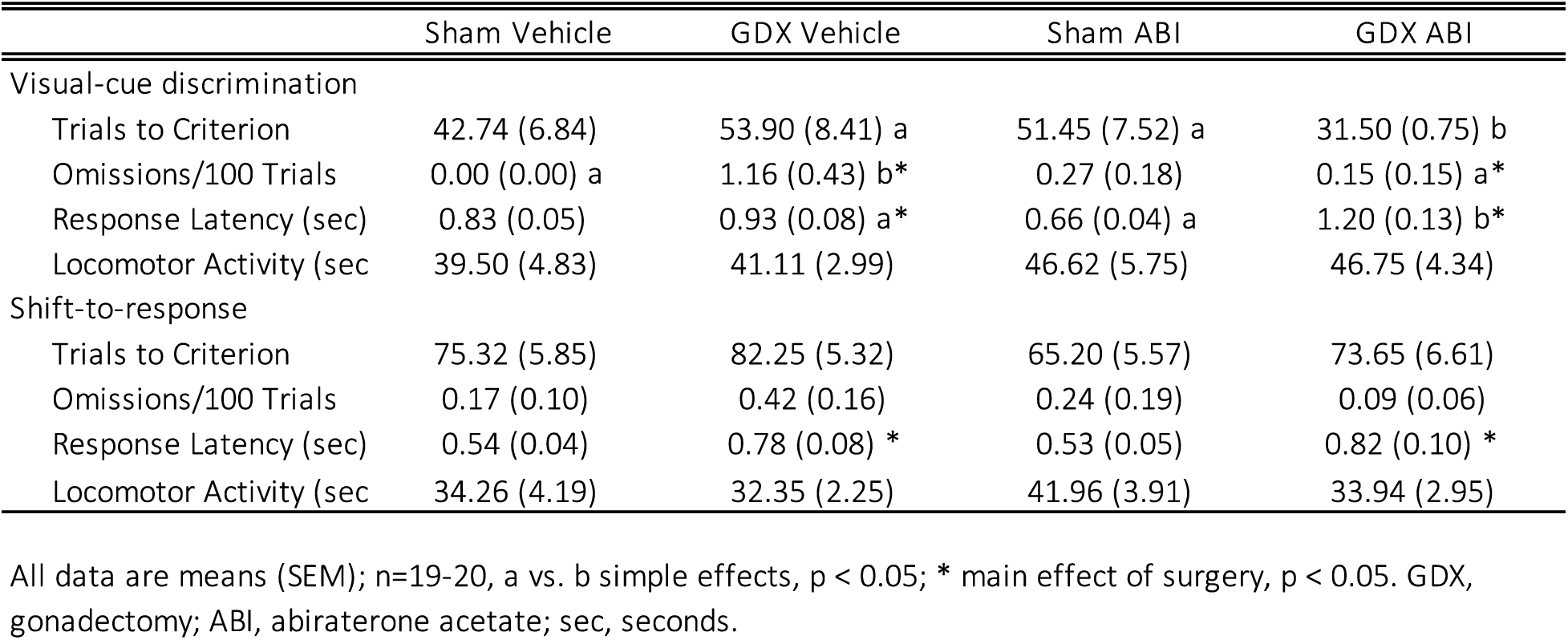
Selected behavioural parameters for strategy set-shifting

We also analyzed omissions, response latencies, and locomotor activity during visual-cue discrimination learning. While omission rates remained low overall, these were affected by Drug and Surgery (Drug: F_1,75_ = 2.27, p = 0.14; Surgery: F_1,75_ = 4.41, p = 0.04; Drug x Surgery: F_1,75_ = 6.62, p = 0.01) on Day 1 (Table 1). Specifically, GDX increased omission rates in Vehicle animals (F_1,75_ = 10.78, p < 0.01), whereas ABI treatment reduced omissions in GDX animals (F_1,75_ = 8.44, p < 0.01) to levels comparable to Sham+Vehicle animals. Drug or Surgery did not affect locomotion (Drug: F_1,75_ = 1.94; p = 0.17; Surgery: F_1,75_ = 0.04; p = 0.85; Drug x Surgery: F_1,75_ = 0.03; p = 0.87). Drug and Surgery affected response latencies (Drug: F_1,75_ = 0.38, p = 0.54; Surgery: F_1,75_ = 15.90, p < 0.01; Drug × Surgery: F_1,75_ = 7.63, p < 0.01). GDX+ABI rats were slower to respond than GDX+Vehicle rats (F_1,75_ = 5.79, p = 0.02) and GDX+ABI rats also had a significantly increased response latency compared to Sham+ABI rats (F_1,75_ = 23.08, p < 0.01; Table 1).

#### Day 2: Shift-to-response discrimination

On the following day, we examined if our manipulations affected recall of the previously acquired visual cue strategy during 20 reminder trials prior to the strategy shift. Analysis of the accuracy data from this part of the session did not yield main effects of Drug or Surgery, but did reveal a significant interaction between the two factors (Drug: F_1,75_ = 0.68, p = 0.41; Surgery: F_1,75_ = 0.30, p = 0.58; Drug × Surgery: F_1,75_ = 5.27, p = 0.02; Figure 2C). This interaction was driven by the fact that the GDX+Vehicle group showed better recall of the visual-cue rule compared to the Sham+Vehicle group (F_1,75_ = 4.00, p = 0.05) and the GDX+ABI group (F_1,75_ = 4.94, p = 0.03; Figure 2C). More importantly, both ABI-treated groups performed similarly to the Sham+Vehicle group suggesting that ABI treatment did not affect the ability to recall the visual-cue rule.

During the shift-to-response discrimination, we found no main effects and no significant interaction on trials to criterion (Drug: F_1,75_ = 2.55, p = 0.11; Surgery: F_1,75_ = 1.72, p = 0.19; Drug × Surgery: F_1,75_ = 0.02, p = 0.90; Table 1). However, on the more sensitive measure of errors to criterion^51, 61–63^, we did observe a significant main effect of Drug (Drug: F_1,75_ = 3.93, p = 0.05). This effect occurred in the absence of either a main effect of Surgery or interaction between the two factors (Surgery: F_1,75_ = 0.58, p = 0.45; Drug × Surgery: F_1,75_ = 0.01, p = 0.91; Figure 2E). Furthermore, the analysis also showed a significant effect on the type of errors that were made (F _2,150_ = 44.08, p < 0.01), even though, this was not affected by the ABI treatment (Drug × Error Type interaction: F_2,150_ = 1.27, p = 0.29). Thus, irrespective of Surgery condition, ABI treated rats made fewer errors to criterion compared to Vehicle treated rats during the shift. Importantly, the GDX+Vehicle group performed similarly to the Sham+Vehicle group. Even though there was no significant Drug x Error type interaction, exploratory analyses showed that ABI treatment tended to reduce perseverative errors (F_1,77_ = 2.87, p = 0.09; One-way ANOVA; Figure 2F) but not regressive errors (F_1,77_ = 0.05, p = 0.82) or never-reinforced errors (F_1,77_ = 1.84, p = 0.18). Thus, the reduction in total errors seems to be driven primarily by a reduction in tendency to persist on using the previously relevant strategy following a change in rule contingencies.

We investigated whether these results could be driven by differences in motivation or activity. Drug and Surgery did not affect omissions (Drug: F_1,75_ = 0.93, p = 0.34; Surgery: F_1,75_ = 0.13, p = 0.72; Drug x Surgery: F_1,75_ = 2.24, p = 0.14) and locomotion (Drug: F_1,75_ = 1.87; p = 0.18; Surgery: F_1,75_ = 2.13; p = 0.15; Drug x Surgery: F_1,75_ = 0.81, p = 0.37; Table 1). Surgery increased latency to respond during the shift (Drug: F_1,75_ = 0.03, p = 0.87; Surgery: F_1,75_ = 13.46, p < 0.01; Surgery x Drug: F_1,75_ = 0.14, p = 0.71; Table 1). GDX animals weighed less in this experiment (Drug: F_1,75_ = 0.05, p = 0.82; Surgery: F_1,75_ = 6.90, p = 0.01; Drug × Surgery: F_1,75_ = 0.01, p = 0.94; Table S3). Collectively, these results show that ABI tends to improve shifting between strategies.

### Steroids

#### Testosterone

T is an important male sex hormone and 95% of it are produced constantly *de novo* by the Leydig cells in the testes^64^. T is released into the circulation and exerts its biological functions by binding to ARs in various target tissues. GDX and ABI treatment, alone or in combination, were able to significantly and effectively eliminate T in whole blood (Drug: F_1,35_ = 16.02, p < 0.001; Surgery: F_1,35_ = 16.02, p < 0.001; Surgery x Drug: F_1,35_ = 16.02, p < 0.001; Figure 3A). In detail, we could detect T at normal levels only in Sham+Vehicle animals and both, GDX and ABI, significantly reduced T levels in whole blood (p < 0.001).

**Figure 3.**
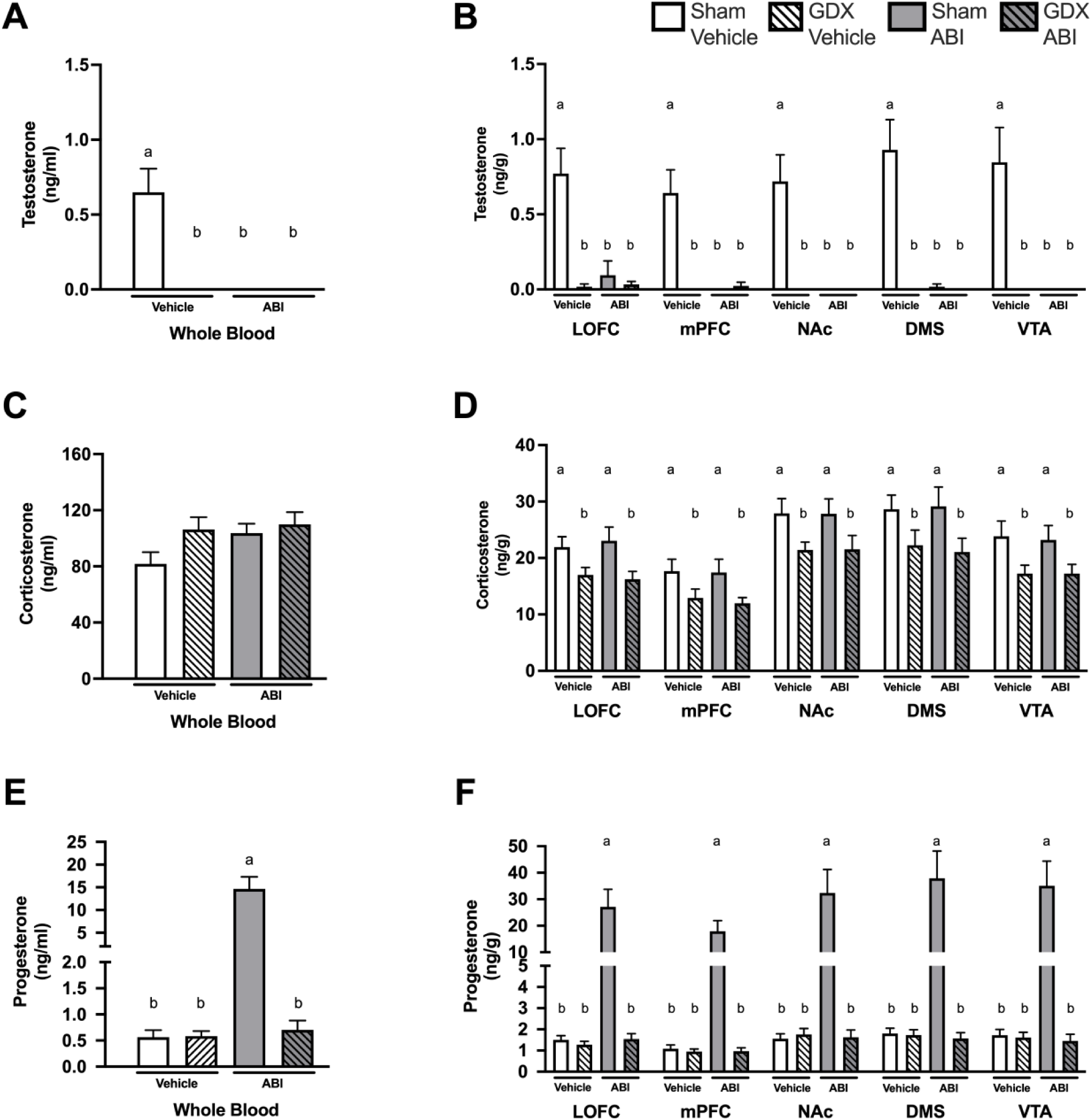
Steroid levels in the strategy set-shifting task: Experiment 1. **A** T is present in whole blood in Sham+Vehicle rats. GDX and ABI treatment greatly reduce T levels in whole blood. **B** T can be found in the brain of Sham Vehicle rats but not in rats treated with ABI or rats that underwent GDX. **C** GDX resulted in a trend to increase levels of corticosterone in whole blood of Vehicle and ABI treated animals. **D** GDX led to a significant reduction of corticosterone in different brain regions of Vehicle and ABI treated rats. **E** ABI treatment in Sham animals led to a significant increase of progesterone in whole blood. **F** ABI treatment in Sham animals led to a significant increase of progesterone in all examined brain regions. ABI, abiraterone acetate; GDX, gonadectomy; mPFC, medial prefrontal cortex; NAc, nucleus accumbens; DMS, dorsomedial striatum; VTA, ventral tegmental area. n=10. Data are presented as mean + SEM. A, B, E, F, **a** and **b** indicate simple effects. For panel D, **a** and **b** indicate main effect of surgery.

Brain T levels were highest in Sham+Vehicle animals and not detectable in most Sham+ABI, GDX+Vehicle, and GDX+ABI animals (Drug: F_1,35-36_ ≥ 11.48, p ≤ 0.002; Surgery: F_1,35-36_ ≥ 13.28, *p* < 0.001; Drug × Surgery: F_1,35-36_ > 12.59, *p* < 0.001; Figure 3B). ABI treatment reduced T levels in all brain regions in Sham animals (F_1,35-36_ > 24.05, p < 0.001) and GDX reduced T in Vehicle animals (F_1,35-36_ > 26.55, p < 0.001; Figure 3B). Sham+Vehicle rats had higher T than GDX+ABI rats (F_1,35-36_ > 26.55, p < 0.001).

#### Corticosterone

Irrespective of drug condition, GDX animals tended towards having higher levels of CORT compared to Sham animals in whole blood (Drug: F_1,35_ = 2.38, p = 0.13; Surgery: F_1,35_ = 3.43, p = 0.07; Drug x Surgery: F_1,35_ = 1.20, p = 0.28; Figure 3C).

In contrast to our observations in whole blood, GDX animals had lower levels of CORT compared to Sham animals across all brain regions, irrespective of ABI treatment (Drug: F_1,35-36_ < 0.11, p ≥ 0.75; Surgery: F_1,35-36_ > 6.75, p ≤ 0.01; Drug × Surgery: F_1,35-36_ < 0.28, p > 0.60; Figure 3D). There was no effect of ABI on CORT.

#### Progesterone

Drug and Surgery affected progesterone levels in whole blood (Drug: F_1,35_ = 33.47, p < 0.01; Surgery: F_1,35_ = 32.16, p < 0.001; Drug × Surgery: F_1,35_ = 32.35, p < 0.001; Figure 3E). ABI significantly increased progesterone in Sham animals in whole blood (F_1,35_ = 38.45, p < 0.001), whereas GDX prevented this increase in the ABI rats (F_1,35_ = 41.14, p < 0.001). Thus, Sham+ABI subjects had higher circulating progesterone levels compared to all other groups.

Drug and Surgery significantly affected levels of progesterone in the brain (Drug: F_1,35-36_ > 11.40, p < 0.002; Surgery: F_1,35-36_ > 11.30, p < 0.002; Drug × Surgery: F_1,35-36_ > 11.60, p < 0.002; Figure 3F). In all brain regions, ABI increased progesterone in Sham animals (F_1, 35-36_ > 23.64, p < 0.001) and GDX counteracted the ABI-induced increase in progesterone levels (F_1,35-36_ > 22.29, p < 0.001). Sham+ABI subjects had the highest progesterone levels compared to all other groups across all brain regions.

### 17-beta-estradiol and DHEA

*17-beta-*estradiol and DHEA were not detectable in any samples.

### Tyrosine hydroxylase histology

In the cortical regions, we saw region-specific effects of GDX and ABI treatment. GDX reduced TH staining in the MOFC (Drug: F_1,35_ = 0.51, p = 0.48; Surgery: F_1,35_ = 9.55, p < 0.01; Drug x Surgery: F_1,35_ = 1.75, p = 0.19; Figure 4A) but not LOFC (Drug: F_1,35_ = 0.01, p = 0.92; Surgery: F_1,35_ = 0.66, p = 0.42; Drug x Surgery: F_1,35_ < 0.01, p = 0.96) and there were no effects of ABI treatment in the LOFC and MOFC. In contrast, in the prelimbic mPFC, ABI treatment increased TH-ir but GDX did not affect TH-ir (Drug: F_1,35_ = 4.80, p = 0.04; Surgery: F_1,35_ = 0.01, p = 0.91; Drug x Surgery: F_1,35_ = 0.07, p = 0.80).

**Figure 4.**
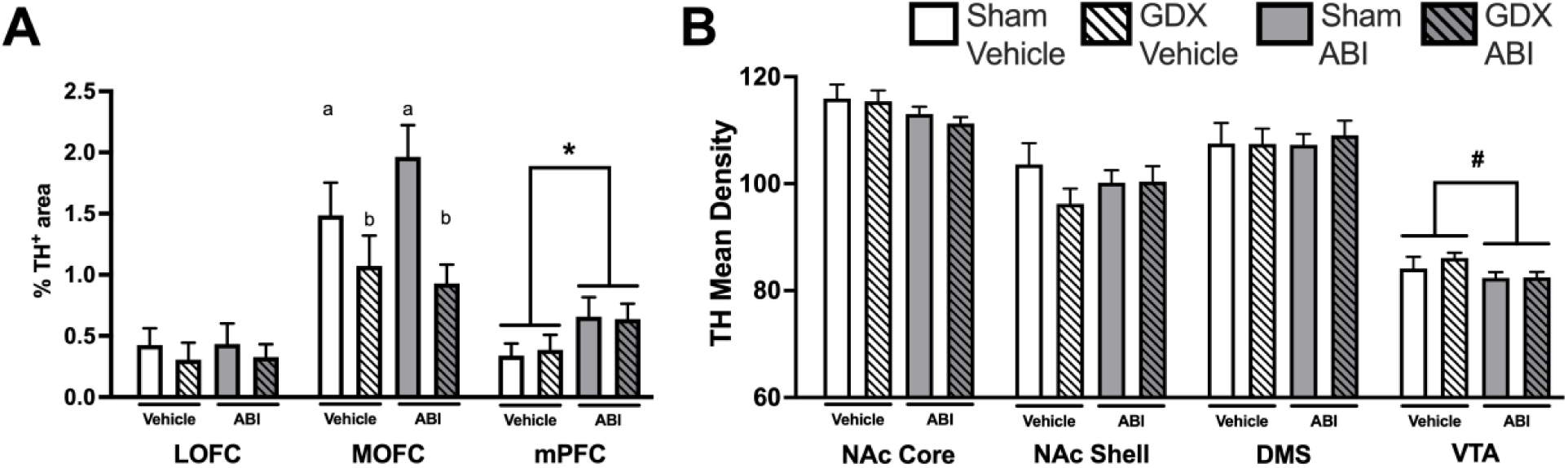
TH-ir in the strategy set-shifting task: Experiment 1. **A** Cortical TH-ir. TH staining expressed as percentage of TH-ir-positive pixels for prefrontal brain regions. GDX led to a reduction in TH staining in the MOFC independent of ABI treatment. Treatment with ABI led to an increase in TH staining in the mPFC independent of GDX. **B** Mesolimbic Tyrosine Hydroxylase. TH staining measured by mean density in the mesocorticolimbic system. GDX did not affec letvels of TH staining in these brain regions but ABI tended to reduce TH-ir in the VTA. Data are presented as mean + SEM. ABI, abiraterone acetate; GDX, gonadectomy; TH-ir, tyrosine hydroxylase immunoreactivity; LOFC, lateral orbitofrontal cortex; MOFC, medial orbitofrontal cortex; mPFC, medial prefrontal cortex; NAc, nucleus accumbens; DMS, dorsomedial striatum; VTA, ventral tegmental area. Data are presented as mean + SEM; n=9-10. *, p < 0.05. **a** and **b** indicate main effect of surgery, p < 0.01, #, p = 0.06.

In the mesolimbic regions, we did not observe effects of GDX or ABI on TH-ir in the NAc (NAcC: Drug: F_1,35_ = 3.60, p = 0.07; Surgery: F_1,35_ = 0.38, p = 0.54; Drug x Surgery: F_1,35_ = 0.11, p = 0.74; NAcS: Drug: F_1,35_ = 0.02, p = 0.90; Surgery: F_1,35_ = 1.41, p = 0.24; Drug x Surgery: F_1,35_ = 1.58, p = 0.22; Figure 4B) and DMS (Drug: F_1,35_ = 0.06, p = 0.82; Surgery: F_1,35_ = 0.09, p = 0.77; Drug x Surgery: F_1,35_ = 0.11, p = 0.75). However, ABI treatment tended to decrease TH-ir in the VTA (Drug: F_1,28_ = 3.85, p = 0.06; Surgery: F_1,28_ = 0.58, p = 0.45; Drug x Surgery: F_1,28_ = 0.47, p = 0.50).

### Experiment 2: Reversal learning

#### Day 1: Response discrimination

The results of the first experiment revealed that ABI improved shifting between different discrimination strategies. In Experiment 2, we examined whether this manipulation might also improve cognitive flexibility during reversal learning, wherein animals use the same basic strategy but must shift between different stimulus-reward associations. To this end, we tested a new cohort of animals on a response discrimination task that depends on egocentric spatial cues (Figure 1). During the initial response discrimination (Figure 5A) Drug and Surgery did not affect trials to criterion (Drug: F_1,59_ < 0.01, p = 0.98; Surgery: F_1,59_ = 0.02, p = 0.89; Drug x Surgery: F_1,59_ = 0.86, p = 0.36; Table 2) or errors to criterion (Drug: F_1,59_ = 0.46, p = 0.50; Surgery: F_1,59_ = 0.07, p = 0.79; Drug x Surgery: F_1,59_ = 0.18, p = 0.67; Figure 5B). Thus, reducing androgen levels does not facilitate the initial learning of a response discrimination.

**Figure 5.**
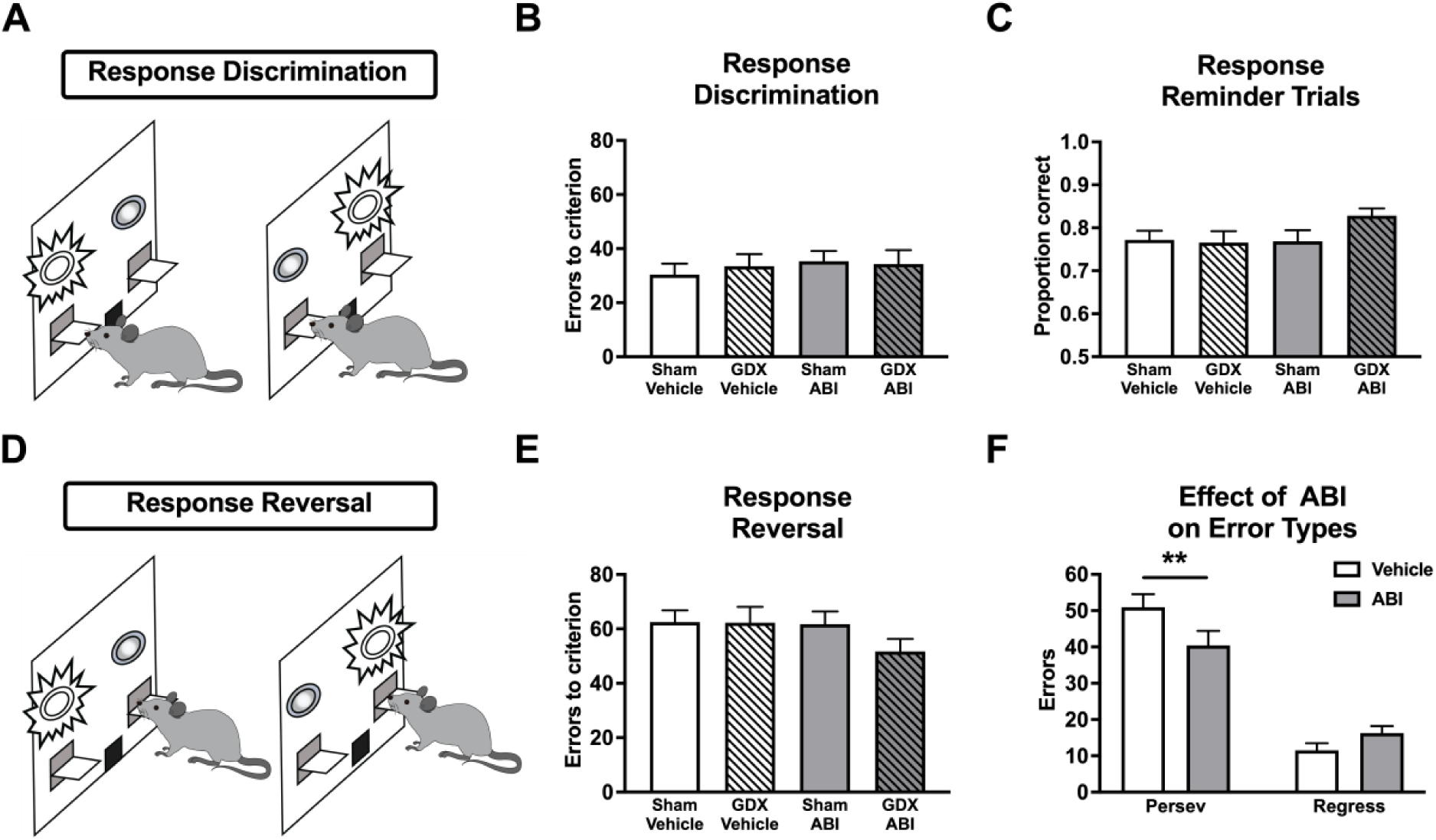
Spatial reversal learning task: Experiment 2. **A** Illustartion of the initial response discrimination. Initially, rats are trained to select the lever opposite of their side bias, irrespective of light position, to receive a reward. **B** Neither GDX nor ABI treatment affected learning of the response discrimination task. **C** All groups showed similar recall of the learned response discrimination. **D** Illustration of the response reversal. Here, rats are shifted to another response discrimination, which requires selection of the opposite lever that was learned during the first response discrimination, irrespective of light position, to receive the reward. **E** Neither GDX nor ABI significantly affected learning the reversal task. **F** ABI treatment (Sham+ABI and GDX+ABI pooled) specifically reduced the number of perseverative errors during reversal learning compared to Vehicle treatment (Sham+Vehicle and GDX+Vehicle pooled). Regressive errors that indicate the ability to maintain the newly learned strategy were comparable between Vehicle and ABI treatment. ABI, Abiraterone acetate; GDX, gonadectomy; persev, perseverative errors; regress, regressive errors. Data are presented as mean + SEM. n=15-16. **, p < 0.01.

**Table 2.** Selected behavioural parameters for reversal learning

|  | Sham Vehicle | GDX Vehicle | Sham ABI | GDX ABI |
| --- | --- | --- | --- | --- |
| Response discrimination |  |  |  |  |
| Trials to Criterion | 76.56 (6.08) | 83.56 (7.46) | 82.50 (5.19) | 77.33 (7.35) |
| Omissions/100 Trials | 0.12 (0.12) | 0.25 (0.12) | 0.00 (0.00) | 0.43 (0.30) |
| Response Latency (sec) | 0.74 (0.05) | 0.78 (0.06) * | 0.56 (0.05) a | 0.93 (0.13) b* |
| Locomotor Activity (sec) | 36.23 (3.07) | 40.04 (2.46) | 38.26 (2.82) | 31.44 (2.73) |
| Response reversal |  |  |  |  |
| Trials to Criterion | 106.81 (6.03) | 105.88 (7.83) | 110.25 (5.01) | 97.80 (6.72) |
| Omissions/100 Trials | 0.09 (0.09) | 0.42 (0.21) | 0.22 (0.18) | 0.35 (0.18) |
| Response Latency (sec) | 0.62 (0.04) | 0.65 (0.06) | 0.49 (0.03) | 0.64 (0.08) |
| Locomotor Activity (sec) | 32.87 (2.99) | 35.08 (2.61) | 33.06 (2.40) | 27.59 (1.87) |

All data are means (SEM); n=15-16; a vs. b simple effects, $p < 0.05$ ; \* main effect of surgery, $p < 0.05$ . GDX, gonadectomy; ABI, abiraterone acetate; sec, seconds.

We examined motivation and activity during the initial response discrimination. Surgery but not Drug affect response latencies (Drug: F_1,59_ = 0.04, p = 0.81; Surgery: F_1,59_ = 7.38, p < 0.01; Drug × Surgery: F_1,59_ = 4.65, p = 0.04). Sham animals responded faster compared to GDX animals. Furthermore, Sham+ABI rats responded faster compared to GDX+ABI rats (F_1,59_ = 11.68, p = 0.001; Table 2). Neither Drug nor Surgery affected omissions (Drug: F_1,59_ = 0.06; p = 0.82; Surgery: F_1,59_ = 2.78; p = 0.10; Drug × Surgery: F_1,59_ = 0.89; p = 0.35) or locomotor activity (Drug: F_1,59_ = 1.39; p = 0.24; Surgery: F_1,59_ = 0.29; p = 0.59; Drug × Surgery: F_1,59_ = 3.65; p = 0.06; Table 2).

#### Day 2: Response reversal

On the following day, rats received 20 reminder trials requiring them to respond on the discrimination they learned on Day 1 (eg; always press the left lever) before experiencing a reversal. We found no differences between groups in the reminder trials (Drug: F_1,59_ = 1.70, p = 0.20; Surgery: F_1,59_ = 1.37, p = 0.25; Drug × Surgery: F_1,59_ = 2.07, p = 0.16; Figure 5C). During the reversal, Drug and Surgery did not affect trials to criterion (Drug: F_1,59_ = 0.13, p = 0.72; Surgery: F_1,59_ = 1.07, p = 0.31; Drug × Surgery: F_1,59_ = 0.79, p = 0.38; Table 2). However, analysis of the different types of errors is a more informative measure of performance, so we employed a mixed-model ANOVA design with Drug and Surgery as the between-subjects factors and Error Type (perseverative, regressive) as the within-subjects factor. This analysis revealed no main effects of Drug (F_1,59_ = 1.31, p = 0.26) or Surgery (F_1,59_ = 2.70, p = 0.11; Figure 5E). However, there was a significant effect of Error Type (F_1,59_ = 72.87, p < 0.001; Figure 5F) for the within-subjects factor, and there was a significant Drug × Error Type interaction (F_1,59_ = 5.53, p = 0.02; Figure 5F). Subsequent analysis of the interaction looking at the effect of Drug for each error type found a significant effect of Drug on perseverative errors (F_1,61_ = 4.18, p = 0.05) but no effect of Drug on regressive errors (F_1,61_ = 3.12, p = 0.08). Thus, ABI-treated subjects made fewer perseverative errors compared to Vehicle-treated subjects. Importantly, GDX+Vehicle animals made a similar number of perseverative errors compared to Sham+Vehicle animals.

To investigate whether these results were associated with changes in motivation or activity levels we analyzed behavioural control measures. Neither Drug nor Surgery affected response latency (Drug: F_1,59_ = 1.65, p = 0.20; Surgery: F_1,59_ = 2.35, p = 0.13; Drug x Surgery: F_1,59_ = 1.21, p = 0.28), omissions (Drug: F_1,59_ = 0.02, p = 0.88; Surgery: F_1,59_ = 1.89, p = 0.17; Drug x Surgery: F_1,59_ = 0.34, p = 0.56), and locomotor activity (Drug: F_1,59_ = 2.11, p = 0.15; Surgery: F_1,59_ = 0.42, p = 0.52; Drug x Surgery: F_1,59_ = 2.32, p = 0.13; Table 2).

As in Experiment 1, GDX but not ABI treatment had a significant effect on body mass (Drug: F_1,59_ = 0.04, p = 0.85; Surgery: F_1,59_ = 39.41, p < 0.001; Drug × Surgery: F_1,59_ < 0.01, p = 0.96; Table S3).

## DISCUSSION

The main finding of the present series of experiments is that T regulates executive functioning in contexts requiring behavioural flexibility. In Experiment 1, ABI treatment enhanced behavioural flexibility on a set shifting task through a reduction in total errors to criterion and thereby a faster transition to an egocentric response rule. This decrease in total errors was mainly driven by a reduction in perseverative errors. Notably, the enhancement of behavioural flexibility in the set shifting experiment was accompanied by an increase in TH-ir in the mPFC and a trending reduction in TH-ir in the VTA. These findings were complemented by the results of Experiment 2, where ABI also enhanced behavioural flexibility. ABI treatment but not GDX reduced perseverative errors during reversal learning. Our results suggest that normal androgen tone in the brain may serve to promote behavioural rigidity or persistence to previously acquired rules or behaviours, and that reducing this tone may facilitate transitions to new strategies.

### ABI decreases persistence in set-shifting and reversal learning

In our experiments, we waited 5 weeks after GDX before we started behavioural testing to allow the brain to potentially upregulate local androgen production^28, 44^. In addition, we used ABI, which is a specific and effective CYP17A1 inhibitor that is fully deconjugated by the liver and crosses the blood-brain barrier^42, 43, 65^. Our main objective was to determine whether inhibition of androgen synthesis or GDX increased behavioural flexibility. T reduces behavioural flexibility, however in most previous studies T levels were increased to supraphysiological levels^4, 6, 9^. For example, Wallin and Wood administered a high dose of T to intact male rats and found impairments in set shifting and reversal learning^4^. The only study that reduced T signalling used the antiandrogen cyproterone acetate in male rats and found reduced persistence in a set shifting task^3^. Here, we not only abolished gonadal T but also abolished extra-gonadal T. We did not observe effects of GDX alone on behavioural flexibility, perhaps because we waited 5 weeks after GDX, to allow the brain to upregulate local T production, as we observed before^28^. However, ABI treatment reduced total errors to criterion during the shift to a new rule in the set shifting task. This indicates that blocking T synthesis by ABI alone, independent of GDX, is sufficient to facilitate behavioural flexibility. These findings suggest that physiological levels of T can promote persistence, and that reducing T may allow for more rapid changes in behaviour. Importantly, the null effect of GDX alone on set-shifting, combined with positive effects of ABI in GDX animals suggests that the proposed actions of T on promoting persistence is mediated at least in part by neural steroid synthesis.

Results of Experiment 2 (reversal learning task), where the rules are reversed within the same stimulus dimension, complement the findings of Experiment 1 (set-shifting task), where the rule shifts between stimulus dimensions. In the reversal learning task, ABI treatment reduced perseverative errors. This further supports a role of neurally-produced androgens in promoting persistence. Even though our effect is mild, it is a consistent effect that we observed over different animal batches and different behavioural tasks.

The effects of ABI on behavioural flexibility are unlikely to be attributable to differences in the encoding of the initial discrimination rules because ABI treatment did not impair performance in the reminder trials, suggesting that reduced T levels do not affect consolidation of a previous rule. Furthermore, the general lack of effects of ABI on locomotion, latency and omissions indicate that the effects of ABI on set shifting and reversal learning are unlikely to be related to changes in activity levels or motivation.

### Effects of ABI on steroids

We predicted that brain T synthesis would be upregulated following long-term GDX, to partially compensate for the loss of gonadal steroids and provide T to key neural circuits^44^. Previously, we detected T in the VTA, NAc and mPFC (but not blood) of long-term GDX male rats. Here, we measured steroids in the prelimbic portion of the mPFC since it is important for set-shifting behaviour. We also examined the LOFC, which is relevant for reversal learning^66^. Surprisingly, we did not detect T in the brain of GDX+Vehicle animals, even though we were able to detect brain T in GDX rats before^28^. Previously, we detected T in the VTA, NAC and mPFC of long-term GDX rats (6 weeks post GDX, either fed *ad libitum* or food restricted). The food-restricted rats were fed 3 h before euthanasia. In contrast, in the present studies, animals were food restricted for a longer time and were fed almost 24h before euthanasia. Food restriction can reduce T levels, which is a likely explanation for lack of detectable brain T in the GDX+Vehicle animals here^67, 68^. Nonetheless, it remains likely that GDX+Vehicle animals produced T in the brain (as seen previously) and that ABI inhibited brain T synthesis in GDX+ABI animals. Another explanation for lack of T in our measurements could be the fast turnover of low T levels in the brain to E_2_ by aromatase, which levels were too low to be detected here^69, 70^.

We also examined CORT and progesterone levels. In rats, androgens often inhibit the HPA axis^71^. Here, neither GDX nor ABI affected baseline CORT levels in the blood. Surprisingly, GDX (but not ABI) reduced baseline CORT levels in the brain. Thus, CORT levels are unlikely to explain the effects of ABI on behavioural flexibility. ABI inhibits Cyp17A1, which converts progestogens into androgens. Thus, when ABI is given to Sham animals, progesterone levels rise in the blood and brain. Progesterone levels were similar in the other groups, suggesting that ABI was not affecting behavioural flexibility via progesterone signalling.

### Effects of ABI on TH

We investigated if changes in TH expression could be an underlying mechanism for the observed behavioural changes, as dopamine signaling in the mesocorticolimbic system is critical for behavioural flexibility. AR is expressed in neurons in prefrontal cortical regions, NAc and VTA, and T alters TH expression^14, 72^. Long-term GDX had no effect on TH-ir in the mPFC or VTA. Interestingly, ABI increased TH-ir in the prelimbic region of the mPFC and tended to reduce TH-ir in the VTA, suggesting that this might be a mechanism by which ABI alters behavioural flexibility. Similar results were observed in our previous study, whereby aged male rats, which have lower T levels, also showed an increase in TH-ir in the infralimbic mPFC but a decrease in the VTA^51^. These aged rats also showed better flexibility in the reversal learning task^51, 73^. In contrast, GDX but not ABI decreased TH-ir in the MOFC, potentially by lack of other gonadal steroids than androgens.

Other studies show that dopamine signaling in the mPFC and VTA is sensitive to T manipulations. Locklear et al. reported that GDX increased burst firing in the VTA, which could be reversed by T but not E_2_. Similarly, GDX increased burst firing in prefrontal cells projecting to the VTA in an androgen-dependent manner. The mPFC is very sensitive to changes in T levels by GDX^30, 32, 33, 75^. Long-term GDX (3-4 weeks) increased TH-ir as well as extracellular dopamine levels in the PFC^32, 33^. Moreover, PFC-to-VTA projections have particularly high AR levels. Effects of T on TH expression can be depending on the behavioural task that animals are trained on, indicating an interaction between steroid levels and behavioural experience^76^. Taken together, the data support the idea that enhancements in behavioural flexibility induced by reducing neural T levels may be mediated through changes in prefrontal dopaminergic circuits.

### Conclusion

In summary, ABI but not GDX increased behavioural flexibility, and ABI had behavioural and neural effects in GDX rats. These data suggest that neurally-produced androgens modulate executive functions mediated by the frontal lobes. In both experiments, ABI improved behavioural flexibility, suggesting that T promotes persistence for a learned strategy or rule. Effects of ABI on TH-ir suggest that these behavioural effects might be driven by alterations in dopamine signalling in the mPFC. These data complement previous studies that demonstrate increasing androgens increases perseveration^4, 6, 7, 9^. Future studies will infuse ABI directly into the brain and examine behaviour. Finally, our data suggest that prostate cancer patients receiving ABI might show changes in executive function^77^.

## Acknowledgements

We thank Chunqi Ma, Ravish Sharma, Brandon Forys, and Hans Adomat for help with data acquisition and analysis. This work was supported by a Discovery Grant from the Natural Sciences and Engineering Research Council of Canada (NSERC) to SBF (12R81349), a Project Grant from the Canadian Institutes of Health Research (CIHR) to KKS (169203), and a Canada Foundation for Innovation (CFI) Grant to KKS (32631). Further funding includes the University of British Columbia Four Year Fellowship, Arts Graduate Award, Aboriginal Graduate Fellowship, and a NSERC Graduate Scholarship to RJT.

