## Supplementary data for "Androgen synthesis inhibition but not gonadectomy reduces persistence during strategy set-shifting and reversal learning in male rats"

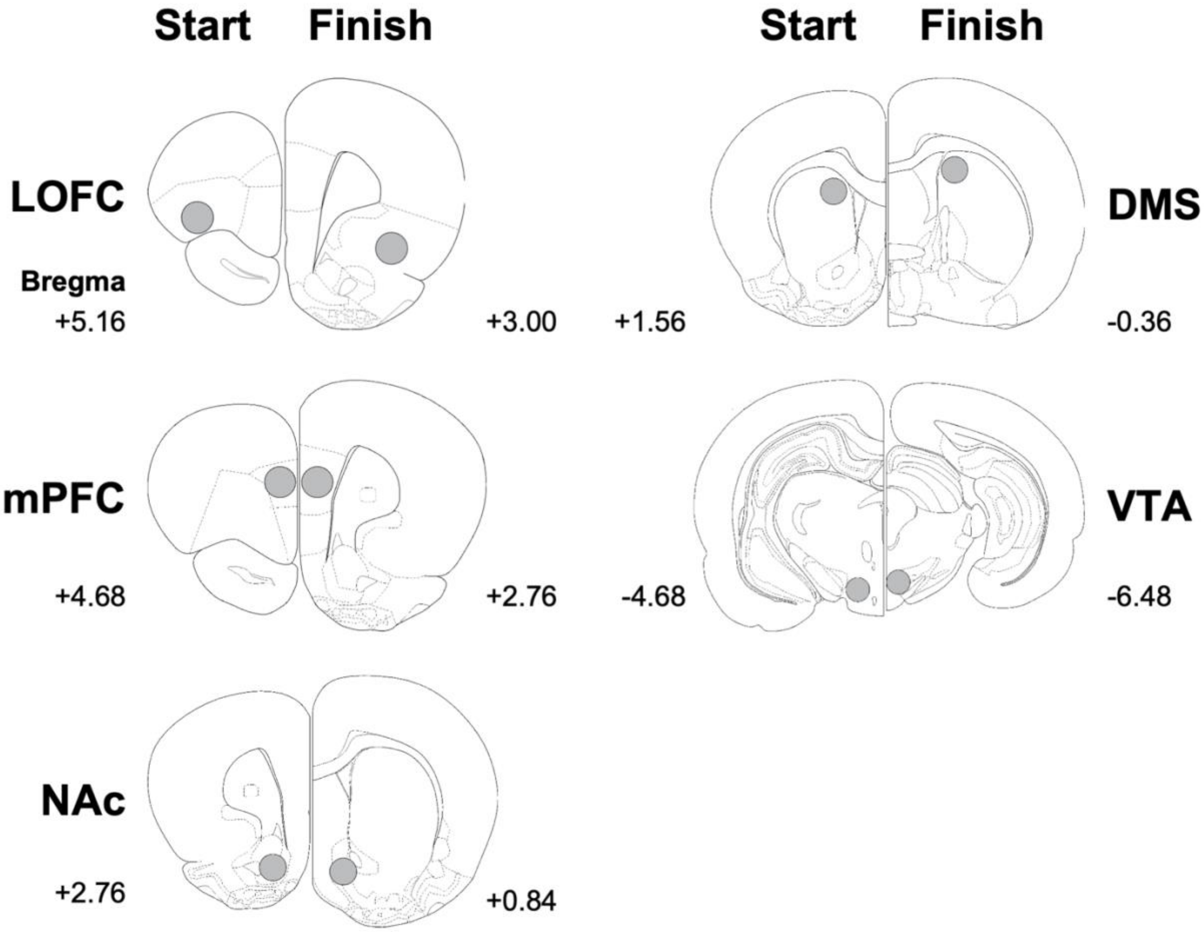


**Figure S1** Schematic illustrating rostral to caudal locations for microdissection performed using 1 mm diameter Palkovits punch for LC-MS/MS steroid analysis. Punch placement was verified by Nissl staining. LOFC, lateral orbitofrontal cortex; MOFC, medial orbitofrontal cortex; mPFC, medial prefrontal cortex; NAc, nucleus accumbens; DMS, dorsomedial striatum; VTA, ventral tegmental area.

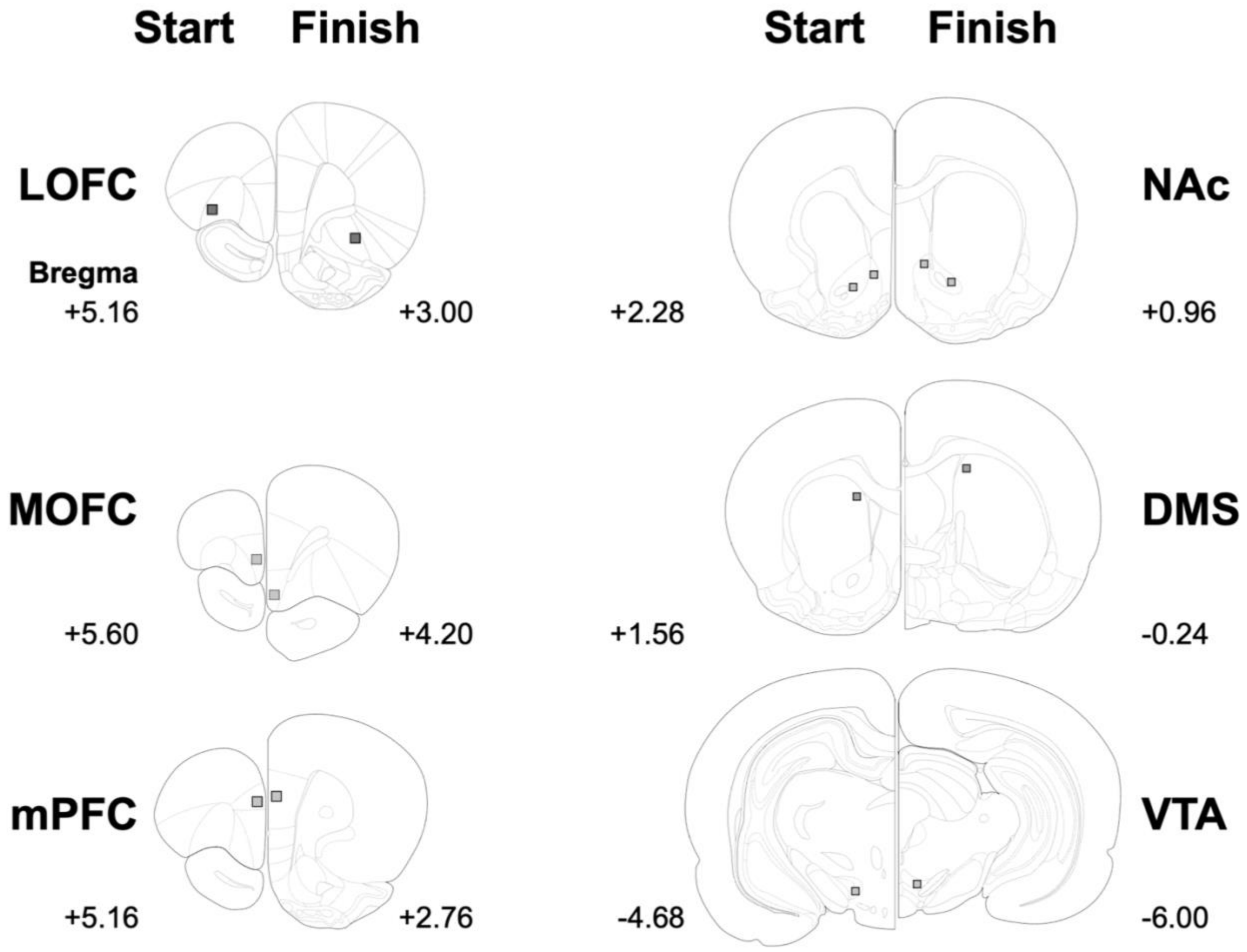


Figure S2 Schematic illustrating rostral to caudal locations for analysis of tyrosine hydroxylase immunoreactivity. LOFC, lateral orbitofrontal cortex; MOFC, medial orbitofrontal cortex; mPFC, medial prefrontal cortex; NAc, nucleus accumbens core and shell; DMS, dorsomedial striatum; VTA, ventral tegmental area.

Table S1. Pilot data: Testosterone levels in serum and brain (ng/ml and ng/g) 4 h after administration of 40 mg/kg ABI p.o. to male rats

| | Vehicle | ABI |
|---|---|---|
| Serum | 0.54 (0.22) | 0.03 (0.01) |
| Brain | | |
| mPFC | 0.45 (0.12) | 0 (0)* |
| VTA | 0.57 (0.17) | 0 (0)* |

All data are means (SEM); n=4; * $p < 0.05$. ABI, abiraterone acetate; mPFC, medial prefrontal cortex; VTA, ventral tegmental area.

Table S2. Pilot data: Abiraterone levels in serum and brain (µg/ml and µg/g) 4 h after administration of 40 mg/kg ABI p.o. in male rats

| | Vehicle | ABI |
|---|---|---|
| Serum | 0 (0) | 0.47 (0.22) |
| Brain | 0 (0) | 1.57 (0.66)* |

All data are means (SEM); n=4; * $p < 0.05$. ABI, abiraterone acetate.

Table S3. Body mass of experimental animals

| | Sham Vehicle | GDX Vehicle | Sham ABI | GDX ABI |
|---|---|---|---|---|
| Experiment 1 | | | | |
| Body Mass (g) | 381.16 (6.00) | 366.35 (5.19) * | 380.30 (6.69) | 364.60 (5.23) * |
| Experiment 2 | | | | |
| Body Mass (g) | 387.00 (4.77) | 358.69 (3.91) * | 385.94 (3.65) | 358.07 (5.46) * |

All data are means (SEM); n=15-20; * main effect of surgery, $p < 0.05$. GDX, gonadectomy; ABI, abiraterone acetate.
